# A horizon for haptic perception

**DOI:** 10.1101/2022.01.19.476921

**Authors:** Luke E. Miller, Felix Jarto, W. Pieter Medendorp

## Abstract

The spatial limits of sensory acquisition (its *sensory horizon*) is a fundamental property of any sensorimotor system. In the present study, we sought to determine the sensory horizon for the human haptic modality. At first blush, it seems obvious that the haptic system is bounded by the space where the body can interact with the environment (e.g., the arm span). However, the human somatosensory system is exquisitely tuned to sensing with tools—blind-cane navigation being a classic example of this. The horizon of haptic perception therefore extends beyond body space, but to what extent is unknown. We first used neuromechanical modelling to determine the theoretical horizon, which we pinpointed as six meters. We then used a psychophysical localization paradigm to behaviorally confirm that humans can haptically localize objects using a six-meter rod. This finding underscores the incredibly flexibility of the brain’s sensorimotor representations, as they can be adapted to sense with an object many times longer than the user’s own body.

## Introduction

Major evolutionary transitions in behavior were often marked by a change in an organism’s sensorimotor horizon. The water-to-land transition in tetrapod evolution increased the sensory volume of vision by nearly one-million-fold (MacIver, Schmitz, Mugan, Murphey, & Mobley, 2017). This led to greater spatial awareness of the environment and is thought to have opened the door for the emergence of long-term planning. The evolution of active sensing systems (e.g., whisking in rodents, electrolocation in weakly electric fish, sonar in cetaceans) afforded the control of perception via the self-driven exchange of energy (and therefore, information) with the environment (Nelson & MacIver, 2006). Here, the sensory horizon often extends to the spatial limit of this energy exchange, such as the tip of a whisker (Bagdasarian et al., 2013) or the field boundaries for electrosensing in weakly electric fish (Babineau, Lewis, & Longtin, 2007). The horizon for sensory acquisition reflects a fundamental property and constraint on any sensorimotor system.

In humans, perhaps the most sophisticated active sensing system involves somatosensory processing during object manipulation (haptic perception; Johansson & Flanagan, 2009; Lederman & Klatzky, 2009), with dedicated processing throughout frontal and parietal cortices (Sathian, 2016). This sophistication was likely driven by one of the most important transitions in human evolution: when we became a tool-making genus three-million years ago (Harmand et al., 2015), which fundamentally reshaped the human hand (Young, 2003) and its neural machinery (Sobinov & Bensmaia, 2021). The volume of sensorimotor interaction (Figure 1A) for our ancestors was no longer limited to arm’s reach (i.e., peripersonal space (Cléry, Guipponi, Wardak, & Hamed, 2015)), but flexibly changed depending upon the tool being used (Maravita & Iriki, 2004). Tool in hand, our ability to haptically perceive objects in the environment now extended beyond the body by at least a meter (Carello, Fitzpatrick, & Turvey, 1992; Miller et al., 2019; Miller et al., 2018; Saig, Gordon, Assa, Arieli, & Ahissar, 2012). The haptic system is exquisitely tuned to interpreting the tool’s location-specific vibratory patterns when contacting an object (Johansson & Flanagan, 2009; Miller et al., 2018), allowing it to function as an extended haptic organ (Miller et al., 2019).

**Figure 1.**
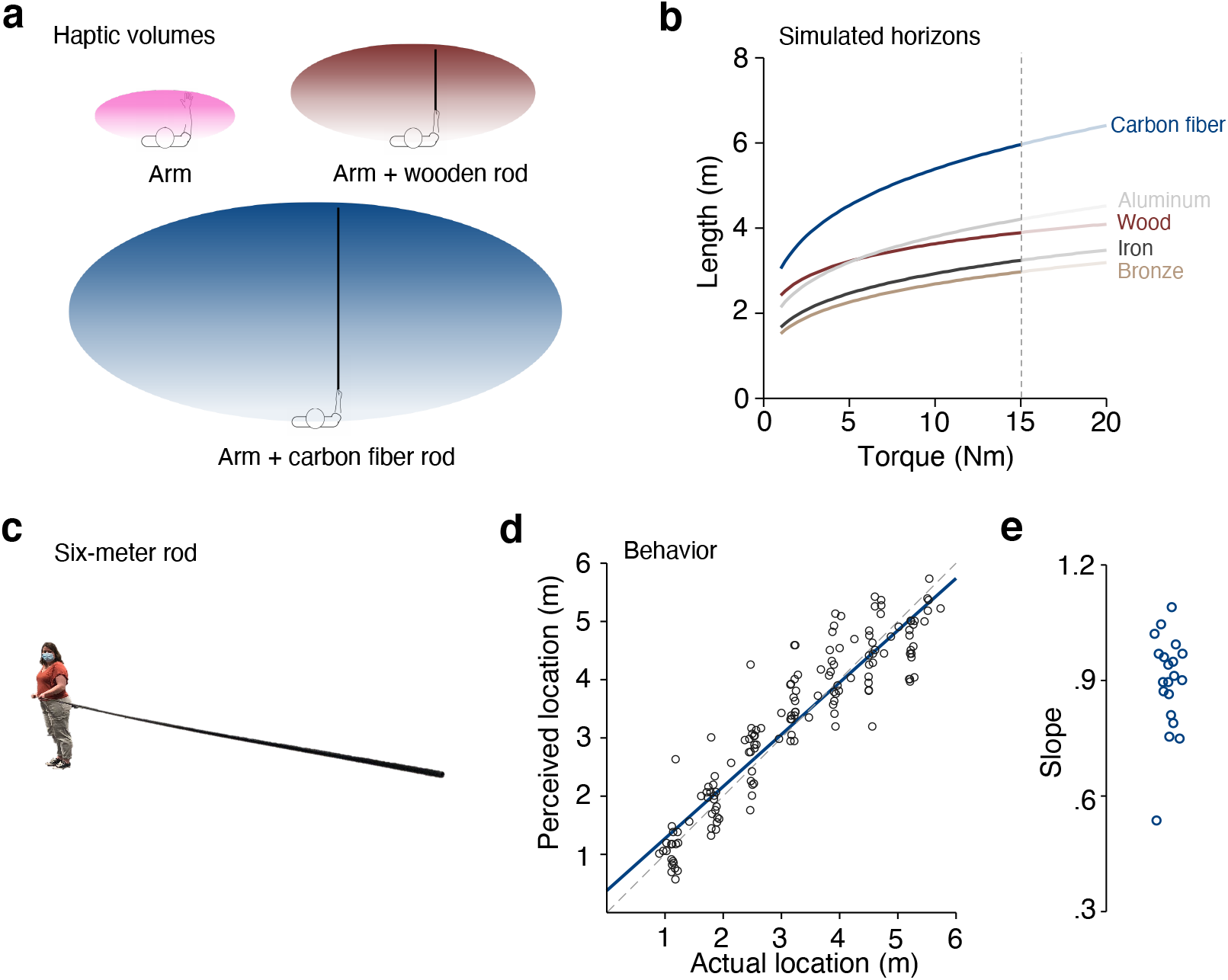
Sensorimotor volumes, simulations, and behavioral results. (**A**) Illustrations of haptic volumes (colored ovals) surrounding the body. Without a tool, the haptic volume is confined to the space that both arms can cover (upper left). With a wooden rod in hand, this volume increases to include the space covered by the length of the rod (upper right). Carbon fiber rods at the theoretical limit expand this volume well beyond the body (bottom). (**B**) Results of neuromechanical simulations for the horizons as a function of torque needed for tool-holding for different materials. The cut-off of fifteen Newton-meter illustrates the likely theoretical limit. Carbon fiber extends the horizon well-beyond the other materials simulated. (**C**) The carbon fiber rod used by participants (n=20) in our behavioral experiment, which was either six (n=6 participants) or 5.5 meters long (n=14 participants) (**D**) The group-level fit of our behavioral experiment (blue line). Each dots corresponds to the perceived location for an individual participant. Note that they are slightly jittered around the actual location for presentation purposes. The dashed grey line corresponds to the identity line. (**E**) The fitted slopes for each participant.

The close link between tools and the human haptic system raises the question of whether or not it has a horizon. Unlike other animals, humans can fabricate the instruments of haptic perception (i.e., tools), expanding their sensorimotor volume ever further away from the body. This is however limited by biomechanical factors (e.g., whether the tool is manipulable), the bandwidth of peripheral receptors (e.g., whether the tool vibrates within their range), and perhaps also by prior sensorimotor experiences with different tool sizes (e.g., it is rare to wield tools longer than five meters). Is there an intrinsic horizon to human haptic perception or does it extend to the limits of any tool that is wieldable?

## Results and Discussion

We first sought to establish a theoretical horizon for the haptic system from first principles. We used neuromechanical modelling to explore the extent to which the haptic horizon is linked to the properties of a tool (here, a rod) that the user could wield—that is, its material (density, elasticity) and geometry (length, diameter). Whether it is possible to use a specific tool to sense the environment depends, in large part, upon the biomechanics and the nervous system of its user. We therefore imposed two major constraints on our simulations, one biomechanical and one neurophysiological.

At the level of biomechanics, a user must be able to lift and transport a rod comfortable if they wish to sense with it. This means that they must be able to apply the necessary torques at the handle in order to keep it at equilibrium and move it from place to place. Our *biomechanical constraint* was that holding the rod stable required no more than fifteen Newton-meters of torque, which we found during piloting to be at the limit of comfortable use. At the level of the nervous system, peripheral receptors must have the necessary bandwidth to pick up the somatosensory information created by tool-object contact. A rod’s resonant frequencies are tightly linked with its material properties and their amplitudes depend upon the location of contact. Our *neurophysiological constraint* was that these vibrations must therefore be within the range of Pacinian receptors (~20–1000 Hz)(Johansson & Flanagan, 2009), which likely encode location information during tool use (Brisben, Hsiao, & Johnson, 1999; Miller et al., 2018).

We simulated the theoretical horizons for various materials that have been important throughout human history (see Methods). The material properties used in our simulations can been seen in Table 1. For a given material, the maximal length of the rod that satisfied both constraints corresponded to its theoretical horizon.

**Table 1.** Material properties used in our simulations.

| Material | Density (kg/m<sup>3</sup>) | Elasticity (GPa) |
| --- | --- | --- |
| Wood | 650 | 14 |
| Bronze | 8000 | 110 |
| Iron | 7800 | 210 |
| Aluminum | 2700 | 68 |
| Carbon fiber | 1600 | 230 |

The results of our simulations (Figure 1B) suggested that the haptic horizon was relatively stable (2-3-meters) throughout most of human material history—neither iron or bronze increased the horizon beyond that of wood. However, the horizon has ballooned with the advent of modern materials. The theoretical horizon of carbon fiber rods was around six meters, providing the greatest expansion of the haptic volume of any material simulated (Figure 1A). However, even if this is the case theoretically, six meters is far removed from most people’s prior tool-wielding experiences. When wielding such a tool, the haptic system may therefore be insensitive for the majority of the length (e.g., after two meters). Alternatively, the sensorimotor system’s internal models may be so well-tuned to the dynamics of tools (Imamizu et al., 2000) that its horizon is adaptable to these edge cases (Miller et al., 2018).

To assess this possibility, we performed a localization task with rods right at theoretical horizon—i.e., satisfying both the biomechanical and neurophysiological constraints. Participants (n=20) wielded carbon fiber rods between 5.5–6 meters (Figure 1C; Table 2) to localize where an object was relative to their body. On a given trial, participants used the rod to strike an object and then verbally reported how far away it was from their hands. To assess the accuracy of localization, we used linear regression to model the perceived distance of the object from the body as a function of the actual distance. The slope value was taken as a measure of localization accuracy.

**Table 2.**
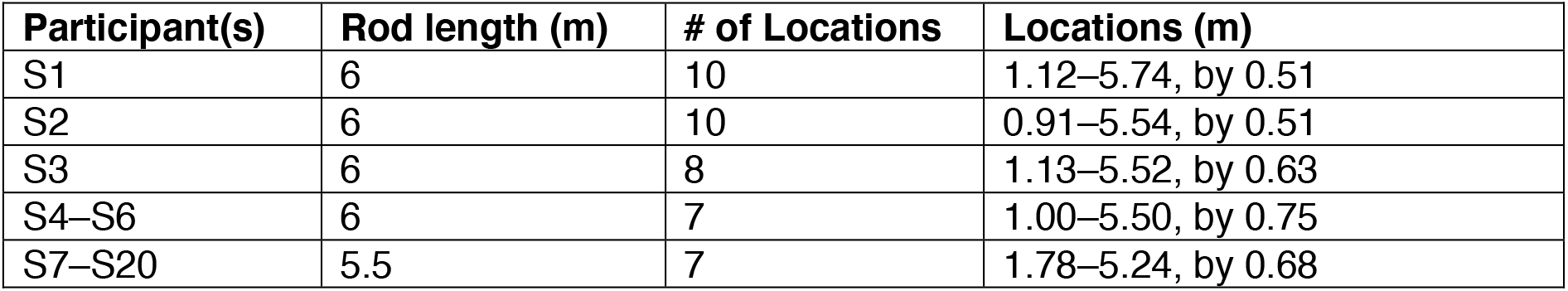
Experimental parameters per participant.

Consistent with what we have observed in prior experiments, performance on the task was near ceiling. Localization in all participants was well fit with a linear model (mean±sem for *R*^*2*^: .91±.02; range: .75–.99). Crucially, participants were highly accurate at localization along the entire length of the rod (Figure 1D), with slopes near identity (slope: 0.89, 95% CI [0.83, 0.95]). This was the case for the majority of our participants (Figure 1E), with only one participant showing slightly lower performance on the task (slope<0.6). The observed range of slope values was consistent with what we have previously measured for rods that were less than one-meter (Miller et al., 2018), despite the fact that length of the rod was >3 times the height of our participants.

Intuitively, our sense of touch and its neural processing (Sathian, 2016) is about events on the body. The proficiency for which humans can extract haptic information from tools— including touch location—calls this intuition into question. Indeed, our results go even further, by demonstrating that the volume of haptic perception actually extends well beyond the physical limits of the body (Figure 1A).

The somatosensory system is highly plastic (Feldman & Brecht, 2005). Its representations are altered by how a limb is used (Kolasinski et al., 2016), change in response to illusions of body size (Tajadura-Jimenez et al., 2012), and are updated when using a tool (Cardinali et al., 2009; Miller, Longo, & Saygin, 2019b) or learning to use an additional robotic thumb (Kieliba, Clode, Maimon-Mor, & Makin, 2021). The current findings further underscore its high degree of flexibility. Even though six meters was outside the range of rods previously wielded by participants, most were still able to use it to accurately localize objects with minimal practice.

This degree of flexibility is likely because users have internalized the relationship between where a rod is contacted and how it vibrates. As discussed above, the frequency content of the vibrations is linked to a rod’s physical properties. For example, the frequencies that a rod resonates at (called *modes*) decrease as a function of its length. The amplitude of each mode instead depends upon where contact occurs. However, unlike the frequencies, these amplitudes are *invariant* across rods and thus provide a universal signal for location (Miller et al., 2018). Given this invariance, it is likely that internal models of tool dynamics (Imamizu et al., 2000). can be readily adapted to be used for different tools, even those with structural properties outside of prior experience.

The proficiency of tool use in humans has led some authors to claim that tools are *embodied*, i.e., treated like extended body parts by the nervous system. Under a structural account of embodiment, tools become ‘incorporated’ with an internal body representation (Maravita & Iriki, 2004), perhaps in multisensory parietal cortices (Iriki, Tanaka, & Iwamura, 1996). However, recent neuroimaging evidence suggests that some neural representations of hands and tools are distinct, even in tool-using experts (Schone, Mor, Baker, & Makin, 2021). This calls the idea of incorporation into question, at least when it comes to the level of neural representation. It is indeed hard to imagine how a neural representation of body space could be so flexible as to incorporate a tool of the length used in the current study, extending its spatial representation out by six meters.

Instead, the present results are more in line with a functional account of embodiment, where sensorimotor computations are merely re-used during tool use. Localizing touch on a rods may involve similar sensorimotor transformations (Yamamoto & Kitazawa, 2001), somatosensory computations (Miller et al., 2022; Miller et al., 2021), and neural processes (Miller et al., 2019) as localizing touch on the body. Given the high localization accuracy observed in the present study, we propose that this is also the case when localizing touch on rods of extreme length. Functional embodiment of extremely long rods may require a radical reconfiguration of how parietal cortex divides up coding for the body and the world (Medendorp & Heed, 2019). We might also expect multisensory neurons coding for peripersonal space (Cléry et al., 2015) to extend their representation to encompass the entire haptic volume, as illustrated in Figure 1A. Future work should utilize edge cases like the present study to fine tune their conceptions of embodiment and its putative neural implementation.

In conclusion, the present study addressed a fundamental question of the human haptic sense: Does it have a horizon and how far beyond the body does it extend? Our results indeed suggest that there is no intrinsic horizon for human haptic perception. Instead, the horizon is inextricably linked to the types of tools available to be wielded. The extension of haptic perception six-meters beyond the body reflects one of the most extreme demonstrations of the flexibility of our sensorimotor system to date.

## Author contributions

L.E.M. and W.P.M conceived of the experiments. L.E.M. performed the neuromechanical modelling. F.J. collected the behavioral data. F.J. and L.E.M. analyzed the behavioral data. L.EM. and W.P.M wrote the paper.

## Data and code availability

All data and code will be made available in repositories upon acceptance of the manuscript.

## Methods

### Neuromechanical modelling

The goal of our neuromechanical modelling was to identify the theoretical sensorimotor horizon of haptic perception from first principles. As discussed in the Main Text, we hypothesized that this horizon is linked to the geometry of a tool that can be used to sense the environment. Whether it is possible to use a specific tool (here, a rod) to sense the environment depends, in large part, upon the biomechanics and the nervous system of its user. We imposed two constraints on our simulations, one biomechanical and one neurophysiological.

To identify the haptic horizon, we simulated the relationship between the physical (material and geometric) properties of a rod and the vibratory information it creates when contacting an object. According to the Euler-Bernoulli theory of beams, striking a rod causes it to resonate at specific frequencies (called, *modes*) whose values are specified by several physical properties: its density, elasticity, cross-sectional radius, and length. Whereas a rod will theoretically resonate at an infinite number of modes, typically only the first five are within the bandwidth of Pacinian mechanoreceptors (20–1000 Hz)(Johansson & Flanagan, 2009). For a specific mode *n* of a rod of uniform material and geometry, its frequency *ω*_*n*_ corresponds to:

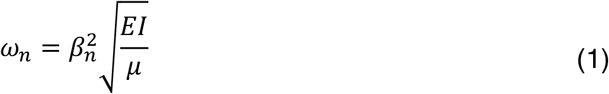

where *E* is the elastic modulus of the rod, *μ* is its mass per unit length, and *I* is its second moment of area. *β*_*n*_ is related to the length of the rod *L*as follows:

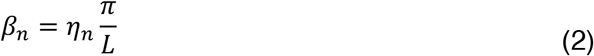

where the value of *β*_*n*_ is determined by solving *cosh*(*β*_*n*_*L*)*cos*(*β*_*n*_*L*) = −1. Longer discussion on these equations can be found in Miller et al., (2018). Importantly, the amplitude of the rod’s resonance in each mode *ω*_*n*_depends upon contact location, a relationship referred to as the mode’s shape. Furthermore, mode shapes are invariant across different rods, making them an ideal cue for contact location so long as the mode frequencies *ω* are detectable by the somatosensory system. We therefore assume that internal models of rod dynamics (Imamizu et al., 2000) are specifically tuned to the shape of the first five modes.

The resonant frequencies of any given rod will relate to its geometry in the following ways: As a rod’s length increases, the frequencies of its modes will decrease until they eventually are outside of the range of mechanoreceptors. Increasing the rod’s radius will increase the frequencies of its modes, but at the expense of increasing the torque needed to hold the rod. This relationship therefore sets up the two constraints we used in our simulations to identify the horizon: (i) the rod cannot require more than 15 Newton-meters of torque to hold; this constraint was identified in pilot experiments. (ii) the frequencies of at least three modes must be within the aforementioned range of Pacinian mechanoreceptors. The longest rod that satisfied both constraints corresponds to the theoretical horizon of haptic perception.

To interrogate this, we simulated how the horizon changed as a function of torque. For a given torque (ranging between 1 and 20 Nm), we found the longest rod whose modes (Equations 1&2) were within the neurophysiological bandwidth. This was done for five historically important materials: wood, bronze, iron, aluminum, and carbon fiber. The densities and elasticities used in our simulations for each material can be found in Table 1. The wooden rod was modelled as a solid rod. All other materials were modelled as pipes with a difference of 2 mm between the inner and outer diameter. The latter was done to maximize the length-torque tradeoff. All simulations were performed in Matlab 2017a (Mathworks).

### Behavioral experiment

#### Participants

Twenty right-handed participants (10 females, 25.75±3.02 years of age) completed the behavioral experiment. All participants had normal or corrected-to-normal vision and no history of neurological impairment. Every participant gave informed consent before the experiment.

#### Experimental setup and paradigm

The goal of our behavioral experiment was to assess how well participants could localize an object using a long carbon fiber rod that was at the theoretical horizon identified in our simulations (see above). The rod was a pipe with an outer diameter of 18 mm and a bore of 16 mm. For six participants, the rod was 6 meters in length (mass: 0.50 kg; torque: 14.72 Nm); for the other fourteen participants it was 5.5 meters in length (mass: 0.46 kg; torque: 12.41 Nm). The object-to-hit was an aluminum bar of ~1.5 meters in length and oriented perpendicular to the rod. This was to ensure that the participant could always strike it successfully. Foam padding was attached on top of the object to reduce the sound of contact. Its height from the floor was adjusted according to the height of the participant. A short three-minute ‘training session’ familiarized the participants with the rod and the object before starting the experiment.

During the task, the participant held the carbon fiber rod with both hands while standing sideways in a comfortable posture (see Figure 1C). Their right hand was positioned at the base of the rod and their left hand was placed 0.5 meters in front. Participants were blindfolded to prevent visual feedback of the rod and the object, and wore noise cancelling headphones that played white noise to prevent auditory feedback of the tool-object contact. In between trials the tip of the rod rested on a cushioned surface to ensure that participants did not have to maintain holding it throughout the entire experiment.

On each trial, the object was placed at one of several locations relative to the body of the participant. Upon receiving a ‘go’ cue (auditory beep), participants lifted the rod and swung it downwards into contact with the object. They then made a verbal numerical judgment about where the object was relative to their body; the value 0 was assigned to the location of their left hand and 100 was assigned to the tip. The experimenter manually recorded their response before continuing to the next trial. The number of object locations and absolute positions varied slightly between participants (see Table 2). For the majority of participants, there were seven locations with eight trials each (56 trials in total). Participants were allowed a break halfway through the experiment.

#### Analysis

We used least-squares linear regression to analyze each participant’s localization accuracy. All verbal judgments were converted into meters. The mean localization judgment was modelled as a function of actual object location. Accuracy was assessed by comparing the group-level confidence intervals around the slope to zero and one. Goodness of fit was quantified using *R*^*2*^.

